# Adaptive loss of shortwave cone opsins during genomic evolution in cartilaginous fish

**DOI:** 10.1101/2024.05.27.594570

**Authors:** Bo Zhang, Yidong Feng, Meiqi Lv, Lei Jia, Yongguan Liao, Xiaoyan Xu, Axel Meyer, Jinsheng Sun, Yumin Li, Yaolei Zhang, Na Zhao, Yunkai Li, Baolong Bao

## Abstract

There is ample evidence of the loss of the shortwave visual protein gene in cartilaginous fish. However, the underlying logic and mechanisms by which organisms need to adapt to the environment behind the opsin loss are still unclear. Here, we report the assembly structure of the whole genome of *Okamejei kenojei* and *Prionace glauca*, identify TcMar difference between skates/rays and sharks/chimaeras, and analyze the distribution characteristics and intra group differentiation of opsin-related genes in cartilaginous fish. By establishing a zebrafish model with short wave opsin gene deletion, we find blue or violet light *via* short-wave sensitive opsins SWS1 or SWS2 can cause the photoreceptor layer thinning (the biomarker for age-related macular degeneration in human eye) through enactive the cell aging. The loss of *sws* is helpful for alleviating the short-wave light damage on the eye. Since the tapetum lucidum in the eye is found broad existence in various cartilaginous fish, a logical hypothesis was creatively proposed that the existence of tapetum lucidum on the retina of cartilaginous fish is interdependent with the loss of shortwave visual protein gene, providing a new perspective and explanatory path for further understanding of the evolution of visual genes in cartilaginous fish.

## Main

The opsins in vertebrates include a rod opsin (RH1) to help see in low light, and four classes of cone opsins (SWS1, SWS2, RH2, and LWS) to see in bright light and detect colors across the visible light spectrum. Throughout evolution, these opsins have been altered, lost, or duplicated in many species to provide unique adaptations for vision underwater. In the agnathan (jawless) lampreys, there is the full complement of one rod and four cone opsin genes^1^, however, short-wave opsin genes *sws*1 and *sws*2 were thought to be lost prior to the divergence of the holocephalan (chimeras) and the elasmobranchs (sharks, skates and rays)^2^. It seems likely that only the RH1, RH2, and LWS opsin genes were retained in ancestral chondrichthyans, all cartilaginous fishes have lost the *sws*1 and *sws*2 opsin genes.

Within the elasmobranchs, most studies showed that the rod-based RH1 opsin and only a single cone opsin has been found in the sharks (Selachii) based on the available molecular and micro spectrophotometric data. Sharks are thought to be cone monochromats without the ability to see colors, whereas rays (Batoidae) are thought to own dichromatic color vision, they have retained both the *rh2* and *lws* cone opsin genes in addition to *rh1*^3–5^. Behavioral experiments in the giant shovelnose ray demonstrate they can discriminate color^6^. The presence of cone pigments and color vision in the other major groups of batoids, the skates and sawfishes, is unknown, although at least two species of skate are thought to possess rod-only retinae^7^.

Since the genome of the elephant shark (*Callorhinchus milii*) was first sequenced, there have several whole genome sequences of cartilaginous fishes available, such as whale shark, brown-banded bamboo shark, cloudy catshark^8–10^, providing the genome level data to outline the elasmobranch opsin evolution. However, additional whole genome sequences, especially the skate genome will be required to provide a more complete picture of elasmobranch opsin evolution. Here we report whole-genome analyses of skate (*Okamejei kenojei*), a cold temperate near bottom dwelling cartilaginous fish in East Asia, and blue shark (*Prionace glauca*), an oceanic species in temperate and tropical waters worldwide, currently the most hunted shark species globally, and combined gene functional analysis to expand capacity for an in-depth explanation of opsin genes evolution in cartilaginous fish.

## Results

### Genome sequencing and assembly

Pacbio continuous long reads (CLR) library for *O. kenojei* and circular consensus sequencing (CCS) libraries for *P. glauca* were constructed, generating 232.59 and 91.22 Gb long reads sequencing data, respectively. Whole genome short-reads and Hi-C sequencing were also carried out for initial contig assemble polishing and chromosome anchoring **(Supplementary Table 1)**. The genomes of *O. kenojei* and *P. glauca* were assembled to be 2.75 and 3.23 Gb, respectively (**Supplementary Table 2**), which are similar to the K-mer estimation results (estimated 2.79Gb and 3.25Gb from 17mer analysis, Supplementary Fig. 1a-b). Contig N50 length of 1.50 and 5.32 Mb (**Supplementary Table 2**), and BUSCO scores of 93.9% and 91.9% (**Supplementary Table 3**) comparable to other published chondrichthyans (**Supplementary Fig. 2**), revealing good assembly continuity and gene completeness. Incorporating the Hi-C data, ∼96.28% (2.65 Gb) and ∼97.85% (3.16 Gb) of the assembled contigs were anchored to 44 and 43 chromosomes, respectively (**Supplementary Fig. 1**c-d). Moreover, 22,965 and 21,229 protein-coding genes were predicted in *O. kenojei* and *P. glauca* genomes with ∼96.75% and ∼98.87% of them functionally annotated based on public databases (**Supplementary Table 4**).

### Transposable elements analysis

More and more evidence proved that repetitive sequences in genomes play critical roles in driving evolution, for example, regulating gene expression^11^. And previous studies have shown that the proportion of repetitive sequences in cartilaginous fish genomes is relatively higher than most teleost^12–14^. Therefore, in this study, we included all available cartilaginous fish genomes published and detected repetitive sequences using the same pipeline to have a deep view of repetitive sequence evolution. Overall, we observed that transposable elements (TEs) accounted for approximately 69.27% of the *O. kenojei* genome and 66.44% of the *P. glauca* genome (**Supplementary Table 5-6**), which do not show significant differences compared to the other cartilaginous fish genomes (**Fig. 1a**). However, when looking into subtypes of DNA transposons, we found that skates (average 3.28%) and rays (2.30%) have significantly higher proportions of DNA/TcMar (Tc1/mariner) elements than sharks (0.14%) and the chimaera (0.03%) (**Fig. 1b**). TcMar is the general name of a series of subcategories, such as TcMar-piggyBac, TcMar-Sleeping Beauty (SB), TcMar-ZB and TcMar-Mos1 and TcMar- Passport, which are widely distributed in vertebrate genomes. These kinds of TEs have been proven to serve as powerful tools for genetic manipulation in zebra fish, elegans, mice, and so on, have an apparent preference for insertion into genes and transcriptional regulatory regions of genes^15–19^, and therefore should contribute greatly to the genome evolution and unique characters of species. One of the most prominent differences between skates/rays and sharks/chimaeras is the body plan, that is, sharks and chimaeras have a body shape like other fish while most skates and rays have flattened bodies and wing-like fins. In our investigated genomes, *Pristis pectinata* is an exception because it also has a body shape more like sharks and the fins are a bit like wings, but not as obvious as those of skate and rays. Intriguingly, the proportion of TcMar (0.33%) in *Pristis pectinata* genome is a little higher than that in sharks/chimaeras, but significantly lower than in skates/rays with wing-like fins. As a consequence, we hypothesize that the burst of TcMar may affect the body plan development of skate and rays by regulating activation of related genes or regulators.

**Fig. 1:**
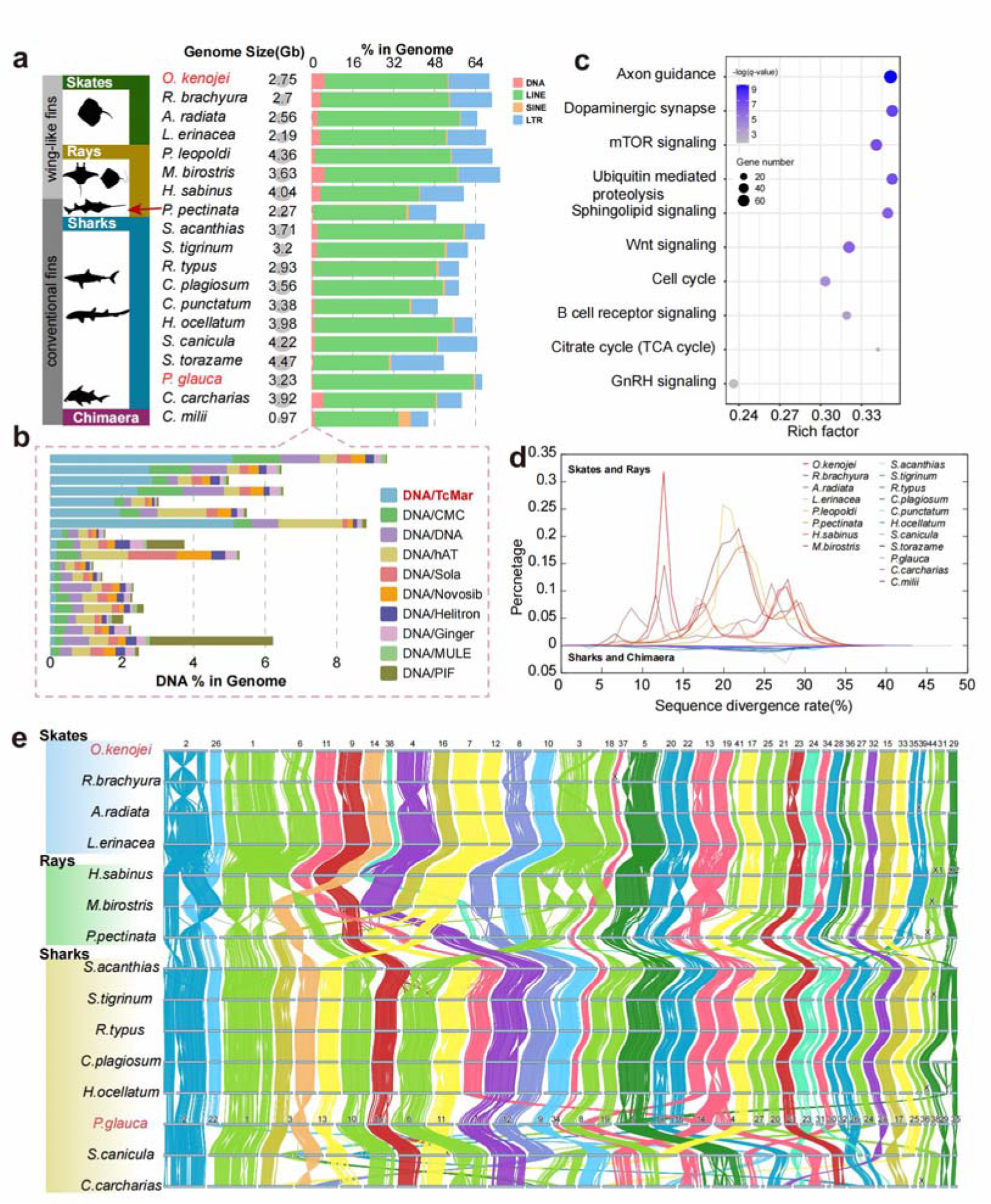
Genomic characteristics of cartilaginous fish. a. Genome sizes and the proportion of DNA transposons, LINEs, SINEs and LTRs in cartilaginous fish genomes, and **b.** the top ten abundant TE super families of cartilaginous fish in DNA transponsons elements. **c**. The top ten KEGG pathways of overlapped TcMar elements blocks in cartilaginous fish genomes included 3,530 genes shared by skates and rays but only 28 genes shared in sharks and chimaeras. **d.** Divergence rates of Tc1/mariner transposable elements in cartilaginous fish genomes. **e.** Pairwise whole-genome alignments across 15 chromosome-level assemblies of cartilaginous fish. Each bar represents a chromosome and the labels above are chromosome IDs.

We next investigated genes whose locus overlapped with TcMar elements blocks in these cartilaginous fish genomes. Interestingly, we identified 3,530 genes shared by skates and rays but only 28 genes shared in sharks and chimaeras. KEGG enrichment analysis showed that the 3,530 genes were significantly (*q*-value < 0.05) enriched in 73 pathways, including Cell cycle, GnRH signaling pathway, Citrate cycle (TCA cycle), B cell receptor signaling pathway, and so on (**Supplementary Data 2**), which are crucial for maintaining life activities. Especially, 60 genes were enriched in Wnt signaling pathway (*q*-value: 1.82×10^-6^), which is a highly conserved system that regulates complex biological processes, for example, controlling axis elongation and cell migration to specific locations thus is critical to the patterning of the entire body plan^20,21^. The 60 genes include most of the members, such as WNT2/3/5/7/9, AXIN1, GSK3B, DVL1, FZD6, LRP6, ROR1/2 and LEF1, which play integral roles in the WNT signaling pathway. The previous study has shown that the WNT/PCP (planar cell polarity) pathway serves as one of the key contributors to skate fin morphology^22^. Therefore, the regulatory expression of genes determines the process of this signaling pathway and TcMar may function in regulating gene expression as they especially expanded in most skate and rays (**Fig. 1c**) and surround these important genes with their regulatory power. Moreover, we also found three rounds of expansion events in skates and at least one round of expansion in rays, which means innovations for these species (**Fig. 1d**)

### Frequent inter-chromosomal rearrangements in cartilaginous fish

The karyotypes vary a lot in cartilaginous fish, which has attracted researchers’ interest^13,22^. However, there are currently no systematic comparisons of chromosomal changes across cartilaginous fishes. Thus, we constructed the syntenic relationships for currently published chromosomal-level assemblies of four skates, three rays, eight sharks and one chimaera to study the changes to show the chromosome rearrangements in cartilaginous fish. We observed an overall conserved pattern of synteny blocks across all genomes. However, there are abundant inter-chromosomal rearrangement events between different orders (**Fig. 1c**). In the superorder Batoidea, the chromosomes between skates maintain the one-to-one correspondence, while in rays, a host of chromosome fusions occurred (chr2 and chr26, chr1 and chr6, chr9 and chr14, chr4 and chr38, chr7 and chr12, chr3 and chr18, chr21 and chr25, chr28 and chr36, chr35 and chr39, chr41 and chr44) in *H. sabinus* and *M. birostris* of Myliobatiformes, leading to the rapid reduction of chromosome numbers of these species. There are 7 chromosomes (chr17, 23, 27, 32, 15, 33 and 29) kept a stable one-to-one chromosome synteny overall in Batoidea, while 15 chromosomes (chr1, 2, 6, 9, 14, 38, 4, 7, 12, 8, 3, 20, 22, 13, and 19) that have experienced more than two fusion and fission events (**Fig. 1c**). On the subject of sharks, *S. acanthias*, with fewer chromosomes, has experienced multiple chromosome fusion events, but has excellent syntenic relationship with the sharks of Orectolobiformes. Conversely, in Carcharhiniformes and Lamniformes, there are frequent chromosomal rearrangement events. Only chr7 and chr34 remain a stable one-to-one collinear relationship among sharks and the chimaera, but 14 chromosomes (chr3, 12, 9, 8, 19, 16, 14, 4, 27, 20, 23, 30, 24 and 28) that have experienced more than two fusion and fission events. In addition, we found that in cartilaginous fish, sex chromosomes may not originate from the same ancestral chromosome, and multiple rearrangement events occur between sex chromosomes and autosomes. These chromosome arrangements may be related to these genome size changes, caused by the burst of repetitive sequence leading to the instability of chromosomes.

To determine the phylogenetic relationships for our newly sequenced species, we first selected 16 representative species, including skates, sharks, and eight teleosts, to conduct gene family analysis. By using SonicParanoid, we identified 10,010 single-copy orthologous genes, based on which, we constructed the phylogenetic tree using the Maximum-likelihood method. The results clearly showed that the *O. kenojei* is unequivocally clustered with *A. radiata*, and the *P. glauca* is the sister branch of *S. torazame*. We also checked the evolutionary relationships of other species used in our analysis, which were consistent with published studies^13^, demonstrating the reliability of our results. We then inferred divergence times with 10 fossil times from the TimeTree database^23^, and found that the skates separated from the sharks about 259 million years ago (**Fig. 2a**), which is also in line with published research^24^. The *O. kenojei* diverged from the *A. radiata* about 33.9 million years ago (MYA), and the *P. glauca* and *S. torazame* diverged from each other about 103.4 MYA.

**Fig. 2:**
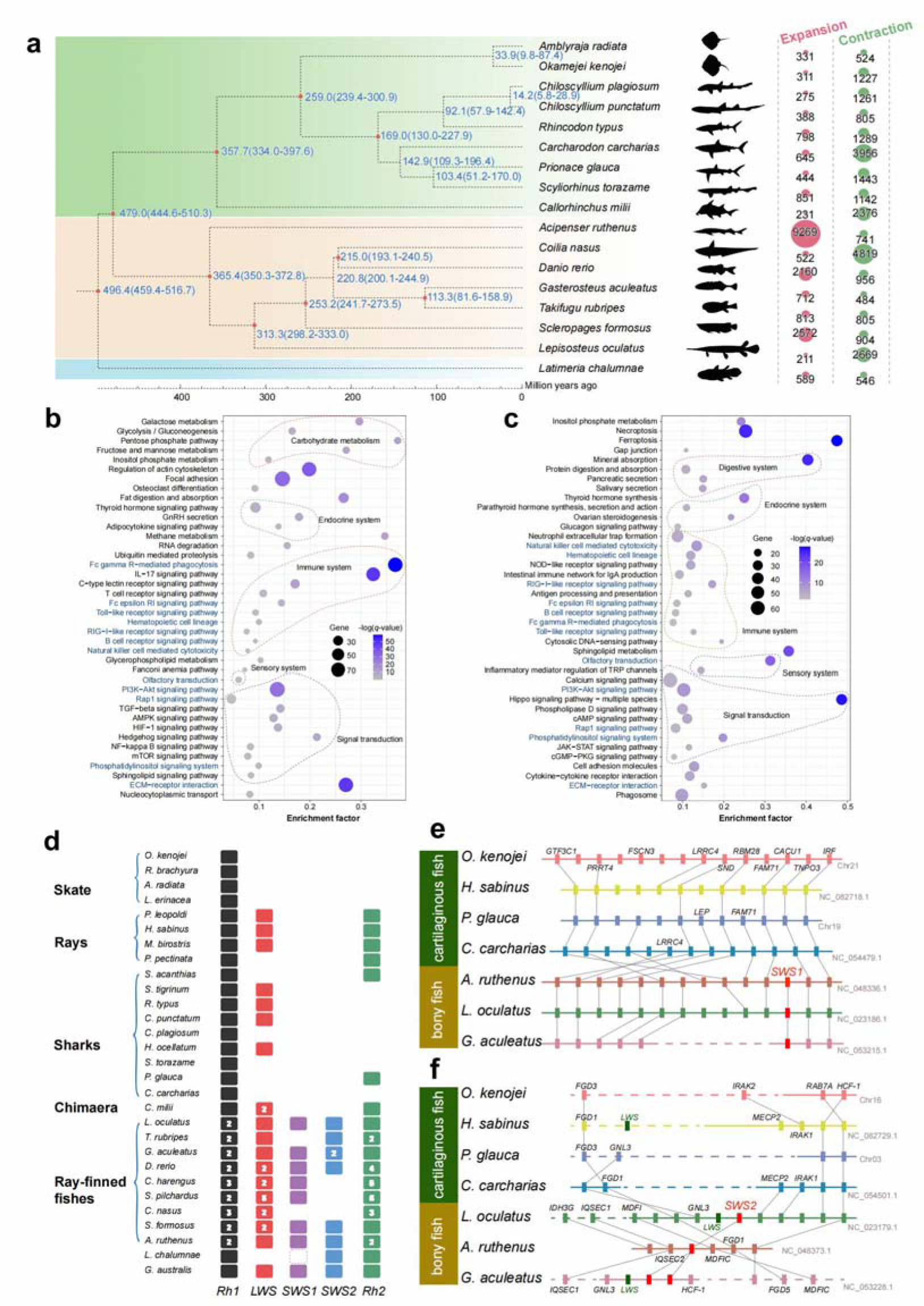
Phylogeny and comparative genomics of cartilaginous fish. **a.** Phylogenetic and gene family expansion and contraction analysis of *O. kenojei* and *P. glauca*. **b,c.** KEGG enrichment analysis of expanded genes in *O. kenojei* and *P. glauca*, respectively, the top 40 enriched pathways are shown. The pathways labeled in blue are shared to both. **d.** Orthology catalogue for opsins of cartilaginous fish. The numbers in the boxes indicate the paralogs numbers. The dashed box represents the pseudogene. **e, f.** The loss of *sws1* and *sws2* genes in cartilaginous fish, respectively. The synteny of genes around *sws1* and *sws2* is illustrated by gray bands connecting orthologs across species.

### Expanded gene families in the two novel genomes

Gene family expansion often makes a species more advantageous for a certain trait. We identified 311 and 444 expanded gene families in *O. kenojei* and *P. glauca* respectively (**Fig. 2a**). In *O. kenojei*, the expanded gene families including 1,246 genes were mainly enriched in pathways relative to carbohydrate metabolism-related pathways, endocrine system, immune system, sensory system and signal transduction (**Fig. 2b**), which are related to survival adaptations. However, as for the *P. glauca*, 444 expanded gene families including 1,611 genes were enriched in pathways quite different in that of *O. kenojei* (**Fig. 2c**) although most of these genes are similarly related to the immune system, endocrine system and signal transduction. Especially, we found that one of the pathways was ovarian steroidogenesis, through which ovarian cells produce hormones for regulation of ovarian function and ovulation^25^. As a viviparous animal, the *P. glauca* has super reproductive capabilities and can produce an average of about 30-50 pups (up to 135 have been recorded) for each breeding, which is much higher than other cartilaginous fishes in the open ocean or even globally^26^. The expanded genes enriched in the pathway included *CYP2J*, *AKR1C3* and *ADCY1*, which played a role in regulatory steroidogenesis^27–29^, and might giving a contribution to multiple births in *P. glauca*.

### Sensory gene repertories

Vertebrates have developed five visual pigment opsin gene lineages including four distinct cone opsins (long-wavelength sensitive, *lws*; short-wavelength sensitive, *sws*1 and *sws*2; middle-wavelength sensitive, *rh*2) originally generated by gene duplication and expressed in cone cells and one rod opsin expressed in rod photoreceptors (rhodopsin, *rh1*)^30^. By checking these genes in cartilaginous fish, we first found that skates lost *lws* and *rh2* while these genes remained in sharks and rays. All of the cartilaginous fish lost *sws*1 and *sws*2 genes which is consistent with previous research^31^ (**Fig. 2d**). However, the loss ways of the two genes seem not the same. For *sws*1, we found good retention of gene blocks in both cartilaginous fish and bony fish (**Fig. 2e**), but this gene was lost separately in cartilaginous fish. However, *sws*2 is more likely to be lost due to drastic changes in changes due to chromosome rearrangements, because compared to bony fish, we cannot find any gene collinearity or blocks in cartilaginous fish (**Fig. 2f**). Maybe cartilaginous fishes have evolved other more advanced sensory systems, and the loss of *sws*1 and *sws*2 is just a use-it-or-lose-it phenomenon or it is that cartilaginous fishes initiatively lost this gene because the existence of these genes directly affects their health? Based on these two hypotheses, we conducted a series of experiments to explore the mechanism of these two gene losses.

### Opsin-dependent short-wave light induced retina injure

What kind of selection pressures leads the universal loss of short-wave sensitive opsins in cartilaginous fish? We hypothesis that there might exist cone opsin-dependent short-wave light induced retina injure in fish. Compared with white light, one-month irradiation of short-wave blue or violet light caused the significant injury of the retina in zebrafish, showing less photoreceptor cells, thinner photoreceptor cell layer (PRL), and thinner pigment epithelial layer (PEL). However, long-wave red or yellow-green light have not caused these injuries (Supplementary Fig.3A&B). In addition, other layers, such as outer nuclear layer (ONL), outer plexus layer (OPL), inner nuclear layer (INL), inner plexus layer (IPL), and nerve fiber layer (NFL) have not injured by the irradiation of short-wave light (Supplementary Fig.3A&B).

Comparative transcriptomic analysis with white light irradiation, the irradiation of shortwave light including blue and violet light upregulated the expression of 54 apoptosis-related genes and 198 cell-aging-related genes (Supplementary Fig.4A), indicating shortwave light might cause cell apoptosis and aging in eyes. Real-time RT-qPCR confirmed the apoptosis marker genes (*caspase3b* and *p21*) and the cell aging marker genes (*mmp65* and *il6*) were upregulated in zebrafish eyes after shortwave irradiation (Supplementary Fig.4B). Immunohistochemistry signal of *caspase3b* and *il6* further confirmed the cell apoptosis and cell aging happen in the eye after shortwave light irradiation (Supplementary Fig.4C). DNA breaks in zebrafish retina were also detected by TUNEL fluorescence staining (Supplementary Fig.4 B&C), which is as previous studies that because of its high energy, blue light induces and accelerates cellular damage *via* the cell apoptosis^32,33^ .

Do the injuries of eye by the short-wave light depend on the short-wave sensitive opsins? In order to understand the role of *sws*, we created *sws1*^-/-^ mutant and *sws2*^-/-^ mutant of zebrafish with CRISPR/Cas9 genomic editing technology (Supplementary Fig.5). In *sws1*^-/-^ mutant and *sws2*^-/-^ mutant of zebrafish after the irradiation of short-wave light (Fig.3A), the photoreceptor cell layer was thicker than that in wild type zebrafish, statistical analysis showed more photoreceptor cells and longer photoreceptor cells in mutant, indicating the injury of photoreceptor cells induced by shortwave light depends short-wave sensitive opsins SWS1 and SWS2. More cell aging genes in the eye were down-regulated in *sws1*^-/-^ mutant and *sws2*^-/-^ mutant after the irradiation of short-wave light (Fig.3B), a total of 157 cellular-aging-related genes were upregulated in *sws2*^-/-^ and 169 cell-aging-related genes were upregulated in *sws1*^-/-^, the relative expression levels of two aging marker genes *p16* and *il6* in the *sws*^-/-^ mutant species were significantly lower than those of the wild type, indicating cell aging might play the major role on the injury of photoreceptor cells. Immunofluorescence of aging marker p16 showed significant low signal in the retina of *sws1*^-/-^ mutant or *sws2*^-/-^ mutant, further confirm the observation of cell aging (Fig.3 C).

**Fig 3:**
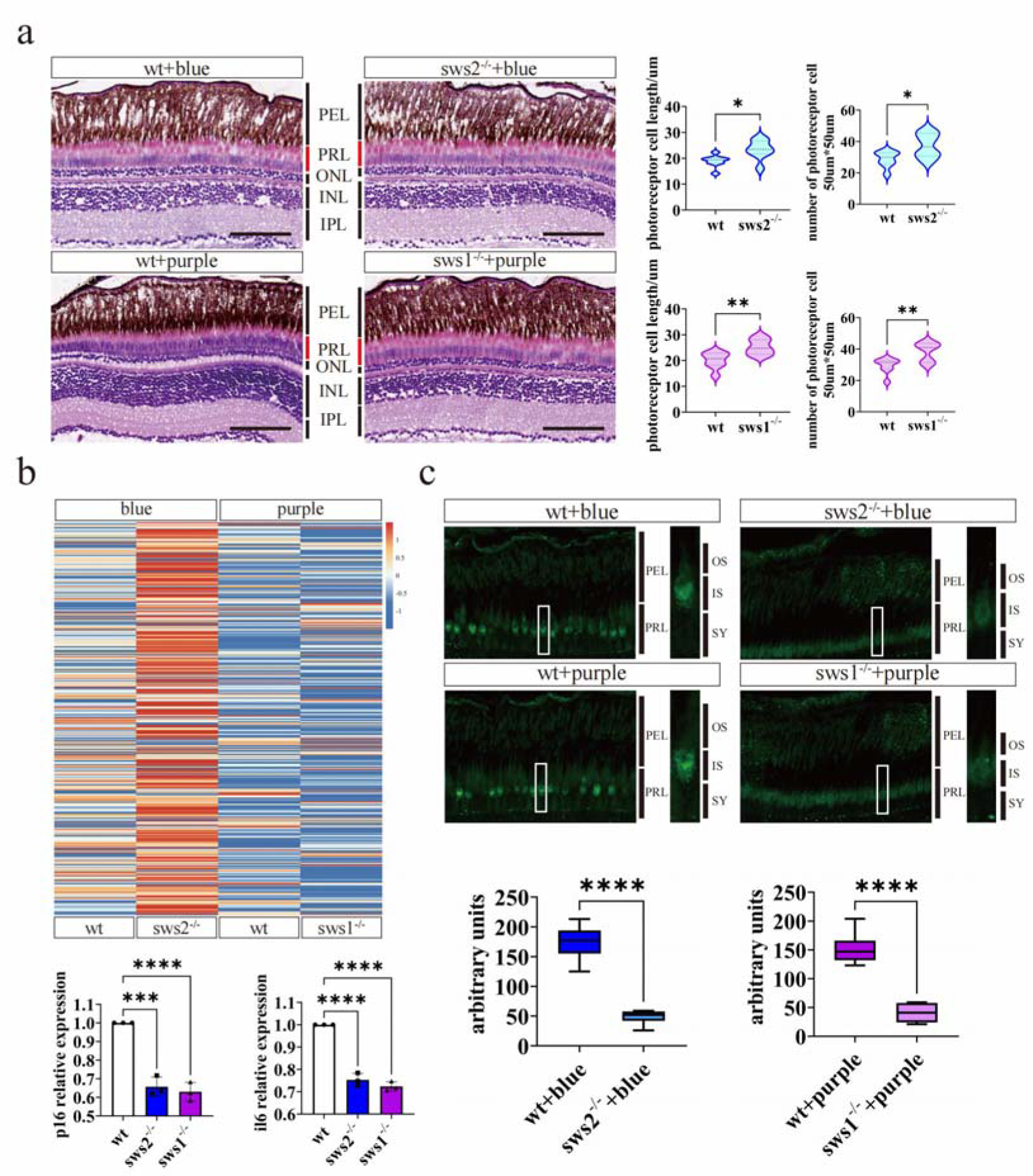
SWS-dependental retina injure and cell aging in zebrafish eyes under short-wave light irradiation. a. Statistical results of the number and length of photoreceptor cells in wild-type zebrafish and *sws2*^-/-^ zebrafish under blue light (λmax=420nm) as well as under purple light (λmax=370nm). n=7, *P<0.05,**P<0.01. **b.** Transcriptomic expression calorigrams of *sws2* mutants and wild-type zebrafish exposed to blue light (λmax=420nm), and *sws1* mutants and wild-type zebrafish exposed to purple light (λmax=370nm). A total of 157 cellular-senescence-related genes were upregulated in *sws2*^-/-^, and 169 cellular-senescence-related genes were upregulated in *sws1*^-/-^. Two aging marker genes, *p16* and *il6*, were verified by qRT-PCR. The relative expression levels of *p16* and *il6* in the two mutant groups were significantly lower than those of the wild type. **c**. The immunofluorescence detection of p16 in retina showed the signal in the inner segment, and the fluorescence intensity of the mutant was significant lower than that of the wild type. n=3. *, P<0.05, **, P<0.01, ***, P<0.001, ****, P<0.0001. PEL:pigment epithelial layer, PRL:photoreceptor layer, ONL:outer nuclear layer, INL:inner nuclear layer, IPL:inner plexiform layer, OS:outer segment, IS:inner segment, SY:synapse.

It is interesting that *sws*-dependent short-wave light can cause the photoreceptor layer thinning through enactive the cell aging, which is the biomarker for age-related macular degeneration (AMD) and the leading cause of vision loss in human being^34^. Indeed, several reports have proposed that the amount of blue light received during an individual’s entire lifespan can be an important factor in the development of AMD^35,36^. In order to further confirm the cell aging depending on zebrafish or human SWS by shortwave light, zebrafish *sws*1, *sws*2 and human *sws* were overexpressed in HEK293 cell line (Supplementary Fig.6). Age-related β-galactosidase staining shows significant signal in *sws*-overexpressing cells, indicating these cells are under aging (Fig.4A). It can be observed the significant signal of p16, cell aging marker protein, in *sws*-overexpressing cells (Fig.4B). These results further show that SWS-dependent short-wave light induced cell aging is be able to happen not only in zebrafish but also in human.

**Fig. 4:**
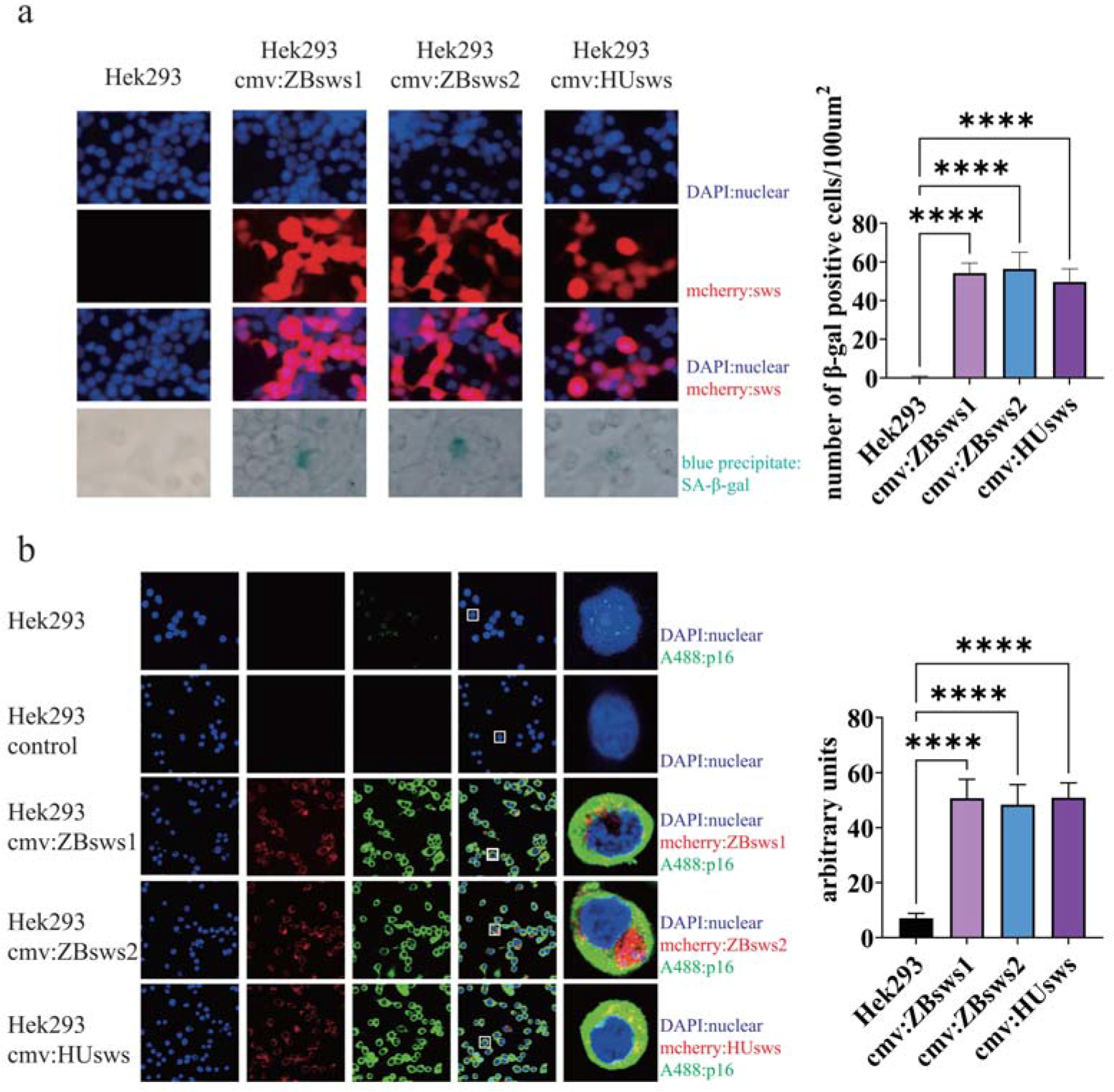
Cell aging detected in *sws*-overexpression cell lines. **a.** Zebrafish *sws1*, *sws2*, and human *sws* overexpressed cell lines exhibited age-related β-galactosidase staining (SA-β-gal; blue precipitate). Untransfected cells did not display SA-β-gal staining. DAPI was used to visualize the nucleus, mcherry for *sws* visualization, and bright-field (BF) imaging for the blue precipitate indicative of SA-β-gal staining. Statistical analysis was conducted on the counts of SA-β-gal positive cells with a sample size of n=100. The level of statistical significance was represented by **** (*P*<0.0001).**b.** Immunofluorescence detection of the p16 under confocal microscopy. HEK293 control served as the negative control group, wherein no primary antibody was added. In the fourth column of the merged image, a white box was used to isolate a single representative cell, which was then enlarged and placed in the fifth column. DAPI was used to stain the nucleus, mcherry for sws visualization, and A488 for p16 labeling. Statistical analysis was conducted on the fluorescence intensity of p16, with a sample size of n=100.

## Discussions

### Two novel genome assemblies

The evolution of cartilaginous fish is of great significance to exploring the evolution of vertebrates. Whole-genome sequencing is an important part of achieving this goal. In this project, we assembled two high-quality chromosomal-level genomes of the cartilaginous fish and discovered a series of important evolutionary events. Firstly, transposable elements are important components of vertebrate genomes, and their diversity and evolution have been widely studied. As a whole, the TE content is higher (45.65-74.79%) in cartilaginous fish genomes than that of the in bony fish (5-56%)^37^.

Although the repeat sequence composition of cartilaginous fish is similar, for example, LINE and LTR are the two main types of repeat sequences, accounting for 86.68%-97.78% of the entire repeats. However, we still found a big difference in subtypes of repeats between skates/rays and sharks/chimaeras, that is the TcMar, which may play a critical role in the body plan. Chromosomal evolutionary events, such as whole-genome duplication and rearrangements, are of great significance to speciation and the shape of new features for species. Based on the two newly assembled and other published genomes, we analyzed the chromosomal rearrangement events of cartilaginous fish found that rearrangement may be an important reason for the loss of genes, for example, *sws2*.

### Evolutionary explanation on the loss of *sws* in cartilaginous fishes

As reported in several published genomes of cartilaginous fishes, the short-wave opsin genes indeed loss in the genomes of skate (*O. kenojei*) and blue shark (*P. glauca*). The phenomenon has aroused extensive attention in the shark research field, it always be discussed in the genomic research of sharks^38–40^ The short-wave opsin genes were thought to be lost prior to the divergence of the holocephalan (chimeras) and the elasmobranchs (sharks, skates and rays)^2^, however, so far there has no reasonable explanation. In this study, we proposed that it exists the injury of short-wave light depending on the short-wave sensitive opsins in fish. And we confirmed by knocking out the gene of short-wave sensitive opsins in zebrafish that blue or violet light *via* short-wave sensitive opsins SWS1 or SWS2 can cause the photoreceptor layer thinning through enactive the cell aging. Moreover, shortwave light indeed induced the cell aging in both zebrafish and human *sws*-overexpressing cells in this study.

Like most of cartilaginous fishes^41–44^, skate (*O. kenojei*) and blue shark (*P. glauca*) have tapetum lucidum that lie in the choroid behind the retina (Fig.5A), the tapetum consists of a single layer of highly reflective, plate-like cells (guanophores), essentially doubling the optical path length of the photoreceptor outer segments and, consequently, the sensitivity of the eye^41^. Therefore, obviously the tapetum lucidum behind the retina will enhance the short-wave sensitive opsins-depending injury on the eyes of cartilaginous fishes (Fig.5B).

**Fig. 5:**
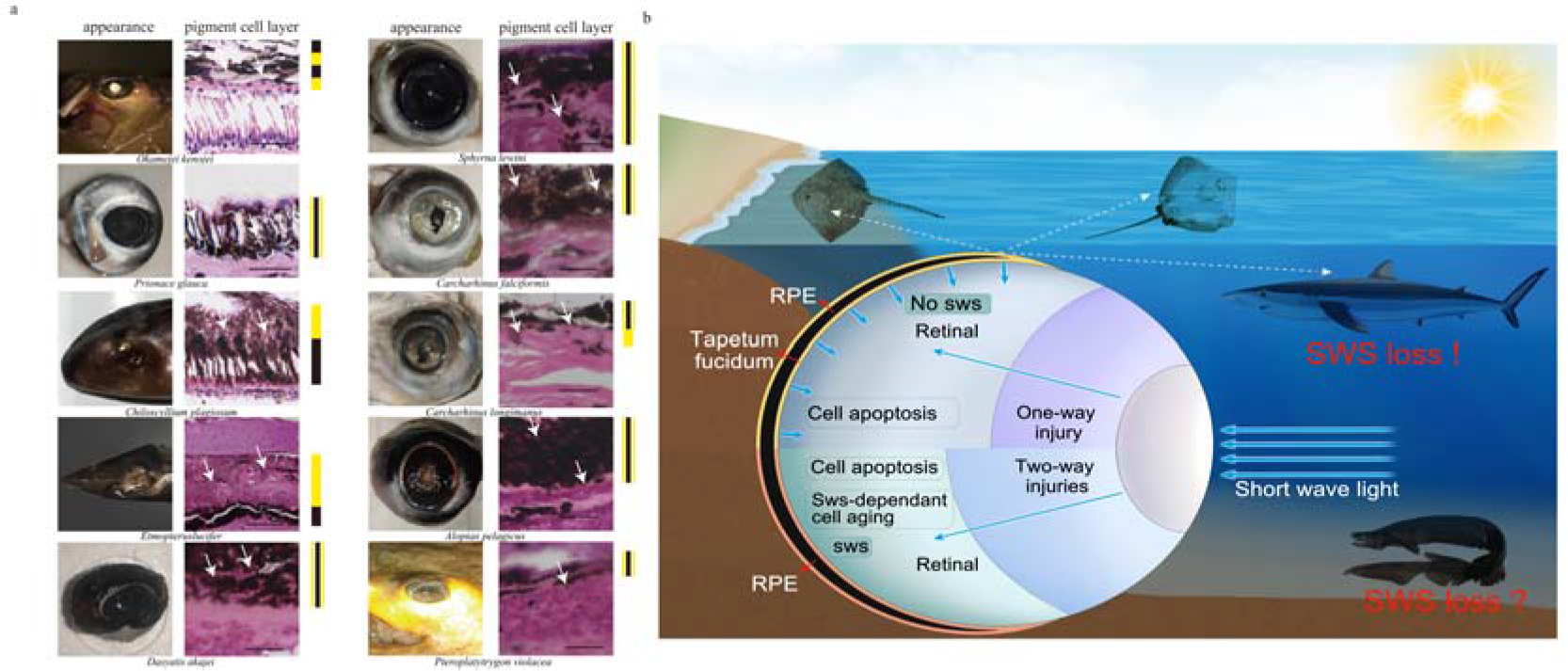
The relationship between tapetum lucidum and sws loss in cartilaginea. a. Tapetum lucidum in the eye of some cartilaginous fish. The appearance was taken with flash in the dark.Tissue sections shows the location of the tapetum lucidum in the pigment cell layer. On the right is a schematic diagram showing the arrangement of melanin (black) and reflective crystals (yellow) in the pigment epithelial cells. **b.** Simplified model to explain cartilaginea sws loss. The lower half of the eye diagram shows two-way injuries to the retina by short-wave light in the presence of sws: apoptosis and sws-dependent cell aging. The upper half of the eye diagram shows that cartilage fish have tapetum lucidum, which is capable of reflecting short waves of light to exacerbate damage to the retina. Ultimately, cartilaginous fish reduce one way of injury by losing sws.

The tapeta do not occur in amphioxus or agnatha, the guanine choroidal tapetum may have arisen in sharks, sturgeon, and lobe-finned fish independent of each other, which might have occurred to allow them to explore deeper depths of the ocean, where light was not as prevalent^45^. When cartilaginous fish species gradually spread into the shallow water area, where light was prevalent, they will face the evolutionary diversity, losing the short-wave sensitive opsins or tapeta, otherwise, the bright light will hurt these fish retinas. When some cartilaginous fishes still live in deep sea, they might be able to own both the short-wave sensitive opsins and the tapeta.

### Age-related macular degeneration (AMD)

Depending the short-wave sensitive opsins, both blue and violet light can cause the photoreceptor layer thinning in zebrafish through enactive the cell aging, which is the biomarker for age-related macular degeneration (AMD) and the leading cause of vision loss in human being^34^. Several reports have proposed that the amount of blue light received during an individual’s entire lifespan can be an important factor in the development AMD^35,36^. In this study, we also provide data that blue light can induce the cell aging of human HEK293T cell line with human *sws*-overexpression, further supporting these postulations on the AMD. Since the AMD is a major health problem in the developed world accounting for approximately half of all blind registrations, and so far there is no AMD-related gene reported, our finding in this study is pretty important to help screening AMD-mark genes on *sws*-dependent transduction pathway and downstream signaling such as cell aging.

### Materials and Methods DNA and RNA sequencing

The *Okamejei kenojei* in this study were caught from Bohai Bay in China. Genomic DNA was extracted from the muscle tissue. One short read paired-end library with an insert size of 350 bp was sequenced using the BGISEQ-500 platform and two 20-kb long read libraries were constructed and sequenced on the Pacific Biosciences RSII platform with the CLR mode using the official standard protocols (Pacific Biosciences). For Hi-C sequencing, the libraries were also constructed and sequenced on BGISEQ-500 platform with 100bp paired-end read using the official standard protocols^46^. RNAs were extracted from six tissues including eye, oarium, dorsal skin, abdomen skin, liver and spermary using the TRIzol kit (Invitrogen, USA) and the libraries were sequenced on the BGISEQ-500 platform (**Supplementary Table 7**). For *Prionace glauca* were from the western Pacific Ocean. A short-read of 350bp library was constructed and sequenced on an Illumina Novaseq platform. For PacBio sequencing, genomic DNA was fragmented to ∼15 kb to construct a long-read library and sequenced on a PacBio Sequel [platform with the CCS mode.

### Genome assembly and chromosome anchoring

The software SMARTdenovo^47^ was used to assemble *O. kenojei* genome using long reads with default parameters. Then the contigs were polished by Pilon (v1.22)^48^ software using the clean short reads filtered by Soapnuke (v1.6.5) with parameters “-M 1 -A 0.4 -d -n 0.02 -l 10 -q 0.1 -Q 2 -G”. Finally, the Hi-C data was mapped to the draft genome using Juicer^49^ and anchored to the chromosome by 3D-DNA pipeline^50^. For the *P. glauca* genome, accurate CCS data were assembled using hifiasm^51^ (v0.16) software. Then purge_dups^52^ (v1.2.6) was used to reduce heterozygous duplications. The chromosomes link using the same methods as that of the *O. kenojei*.

### Genome annotation

For genome repeat annotation, we firstly used RepeatModeler (v1.0.8) and LTR-FINDER^53^ (v1.0.6) to build a custom repeat library and then used RepeatMasker^54^ (v4.0.6) to search the genome repeats against both the custom library and Repbase (v21.01) database, using the parameters ‘-nolow -no_is -norna -engine ncbi’. RepeatProteinMask (v4.0.6) was also used to perform homolog-based search at the protein level with parameters ‘-engine ncbi -noLowSimple -p-value 0.0001’. In addition, tandem repeats were detected by Tandem Repeats Finder^55^ (v4.07) with the parameters ‘-Match 2 -Mismatch 7 -Delta 7 -PM 80 -PI 10 -Minscore 50 -MaxPeriod 2000’. Additionally, repeat contents in other cartilaginous fish for downstream analysis (**Supplementary Table 8**) were detected using the same method above.

For gene annotation, protein sequences of *Chiloscyllium punctatum*, *Scyliorhinus torazame*, *Callorhinchus milii*, *Rhincodon typus*, and *Carcharodon carcharias* (**Supplementary Table 9**) were downloaded from public databases and mapped to the genome using BLAT^56^ software (v35.1), and then gene models were predicted by GeneWise^57^ software (v2.4.1). RNA reads were aligned to the genome using HISAT2^58^, then the transcripts were assembled by Stringtie^59^ (v1.2.2) and the open reading frames were predicted using TransDecoder (v5.5.0) (http://transdecoder.sourceforge.net/). Finally, GLEAN^60^ was used to integrate a non-redundant gene set.

Protein sequences were aligned to different databases for gene function annotation, including Swiss-Prot, TrEMBL^61^, and Kyoto Encyclopedia of Genes and Genomes^62^ (KEGG v105) using BLASTP^63^ (v2.2.26). Function-specific motifs and domains were determined by InterProScan^64^ (v5.60-92.0) according to several protein databases, including Pfam, SMART, PANTHER, PRINTS, PROSITE profiles, and ProSitePatterns and Gene Ontology (GO) annotation results were extracted from the InterProScan results. The protein clustering was performed by Cd-hit^65^ (v4.8.1) with the similarity of 70%.

### Gene family analysis

Gene sequences of *Amblyraja radiata*, *Chiloscyllium plagiosum*, *Callorhinchus milii*, *Carcharodon carcharias*, *Chiloscyllium punctatum*, *Rhincodon typus*, *Scyliorhinus torazame*, *Latimeria chalumnae*, *Lepisosteus oculatus*, *Danio rerio*, *Acipenser ruthenus*, *Coilia nasus*, *Gasterosteus aculeatus*, *Takifugu rubripes*, and *Scleropages Formosus* were downloaded from NCBI database (**Supplementary Table 10**). Then SonicParanoid^66^ was used to identify the orthologous groups among the species and four-fold degenerate (4D) sites were extracted to construct a phylogenetic tree using IQ-TREE^67^ by Maximum Likelihood method. The divergence times were inferred using MCMCtree^68^, and the calibrating fossil times were searched from TimeTree^23^ website: *Latimeria chalumnae* and *Lepisosteus oculatus*, 424-440 Mya; *Latimeria chalumnae* and *Callorhinchus milii*, 442-515 Mya; *Acipenser ruthenus* and *Lepisosteus oculatus*, 345-372 Mya; *Lepisosteus oculatus* and *Danio rerio*, 298-342 Mya; *Scleropages formosus* and *Danio rerio*, 244-301 Mya; *Coilia nasus* and *Danio rerio*,151-246 Mya; *Gasterosteus aculeatus* and *Takifugu rubripes*, 82-174 Mya; *Carcharodon Carcharias* and *Rhincodon typus*, 113-289 Mya; *Callorhinchus milii* and *Amblyraja radiata*, 338-471 Mya; *Chiloscyllium plagiosum* and *Amblyraja radiata*, 245-343 Mya. Then the expansion and contraction of gene families were defined by Café^69^ (v4.0).

### Opsin genes identification

Protein sequences of cartilaginous fish and reference opsin genes of 27 species were downloaded from NCBI and published articles (**Supplementary Table 11**). Then the opsin genes from NCBI databases (**Supplementary Data 1**) were aligned to the whole protein sequences using BLASTP (v2.2.26) with parameters ‘-e 1e-5’. For *Raja brachyura*, *Mobula birostris* and *Squalus acanthias* which were without annotation information, we used miniprot^70^ (v0.12) to align opsin proteins against their genomes to get opsin homologs.

### Zebrafish

The zebrafish utilized in this study comprised the AB wild-type zebrafish, procured from the Shanghai Institute of Biochemistry and Cell Biology, Chinese Academy of Sciences. The wild-type (WT) zebrafish population was maintained under a 14-hour light: 10-hour dark (14 L: 10 D) cycle, and received two daily feedings of *Artemia* nauplii. Embryos were cultivated at a consistent temperature of 27±1°C in egg water, formulated by diluting artificial seawater at a ratio of 1.5:1000 in regular water. The zebrafish broodstock was selectively paired based on a male-to-female ratio of 1:1^71^.

### Light exposure

For violet light exposure, ten *sws*1-mutants (15 dpf) and ten wild-types (15 dpf) zebrafish, separately, were prepared as one group, and three parallel groups were set, placed in a dark room with LED lights on the top, fed twice in the morning and evening, changed the water every day, the irradiation cycle was 14 L: 10 D, and the sustained light stimulation time was 30 days. LED lamp wavelength λ_max_=370 nm, light intensity 2000±100 lx. For blue light exposure, LED lamp wavelength λ_max_=420nm, light intensity 2000±100 lx. The *sws*1-mutants were instead by sws2-mutants, others were same with violet light exposure experiment.

### Preparation of tissue sections

Whole zebrafish are placed in 4% PFA immediately after euthanasia. After an overnight period of fixation, the specimens undergo a series of alcohol washes and dehydration steps, with increasing concentrations of alcohol (75%, 85%, 95%, 100%, every 30 minutes). Subsequently, the samples are rendered transparent using xylene, followed by overnight immersion in paraffin. Finally, 5μm slices were cut from the embedded tissue with a blade, mounted on gelatin-coated slides, and air-dried.

### H&E Staining

The paraffin sections were deparaffinized at 65[and rehydrated using decreasing concentrations of alcohol (100%, 95%, 90%, 80%, 70%, every 2 minutes). Subsequently, they were immersed in hematoxylin dye solution for 5 minutes, followed by rinsing with running water. Differentiation was achieved using a 0.5% hydrochloric acid alcohol solution for 10 seconds, followed by rinsing with running water and immersion in 1% eosin dye solution for 10 seconds. After gradient dehydration, xylene was added to the samples for 1-2 minutes for transparency, and the sections were then mounted with resin.

### Immunohistochemistry

The tissue sections were placed in 3% hydrogen peroxide solution and incubated at room temperature away from light for 25 minutes. The slides were placed in PBS and washed by shaking on a shaking table 3 times. After 5 minutes each time, the serum was sealed, and 3% BSA was added to the tissue in the tissue chemical circle to evenly cover the tissue, and the tissue was sealed at room temperature for 30 minutes. Remove the sealing solution, add PBS to the slices with a certain proportion of primary antibody, and the slices are placed flat in a wet box at 4°C for overnight incubation. The slides were washed in PBS by shaking on a shaker for 3 times, 5 minutes each time. After the slices were slightly dried, the tissue was covered with the secondary antibody (HRP label) of the corresponding species of the primary antibody, and incubated at room temperature for 50 minutes. The slides were placed in PBS and washed by shaking on the decolorizing shaker 3 times, 5min each time. After the sections were slightly dried, the freshly prepared DAB color-developing solution was added to the circle, put in hematoxylin dye solution for 3 minutes, rinsed with running water. After gradient dehydration again, xylene was added to transparent samples for 1-2 min. Finally, the slices were mounted with resin.

### Immunofluorescence and TUNEL

For paraffin sections, the treatment of adding secondary antibodies to drops is the same as immunohistochemistry. After the secondary antibodies were washed, DAPI was added to restain the nucleus and incubated at room temperature for 10 minutes away from light. Finally, the tablets were sealed with anti-fluorescence quenching tablets. For cells grown on the coverslips, cells were fixed with 4% ice-cold paraformaldehyde PBS for 20 minutes and then infiltrated with 0.1% TritonX-100 PBS for 10 minutes. After washing with PBS twice, cells were blocked at 37[for 30 min with 5% BSA, and incubated at 4[overnight with primary antibody. On the second day, the cells were washed with PBS, and then incubated with the corresponding secondary antibody at 37 ° C for 30 minutes, DAPI was re-stained with the nucleus, and the tablets were sealed with anti-fluorescence quenching tablets. For TUNEL staining, tissue slides were subjected to the TUNEL BrightGreen Apoptosis Detection Kit (A112, Vazyme, China). All fluorescence imaging of the sections was conducted using Leica SP8 confocal microscopy. ImageJ software was employed for the statistical analysis of fluorescence intensity, enabling the comparison of mean values between the control and experimental groups.

### Total RNA extraction and cDNA synthesis from eyes

Three samples of zebrafish eye tissue were collected, and RNA extraction was performed following the protocol outlined in the RNA isoPlus specification (9108, TAKARA, Japan). Subsequently, the first strand of cDNA was synthesized using the HiSlid cDNA Synthesis Kit (MKG840, MIKX, China) as per the manufacturer’s instructions.

### RNA-sequencing analysis

The study included eye tissue samples from the following groups: normal feeding wild type (comprising 3 mixed samples of fish eye tissue), blue light-exposed wild type (containing 3 mixed samples of fish eye tissue), violet light-exposed wild type (comprising 3 mixed samples of fish eye tissue), blue light-exposed *sws2*^-/-^ mutant (containing 3 mixed samples of fish eye tissue), and violet light-exposed *sws1*^-/-^ mutant (counting 3 mixed samples of fish eye tissue). Sequencing was conducted by Shanghai Ouyi Biology. Differential expression analysis was performed using the DESeq2.Q value < 0.05 and foldchange > 2 or foldchange < 0.5 was set as the threshold for significantly differential expression gene.

### qRT-PCR

The Real-time PCR analyses were conducted using the Pro SYBR qPCR Mix (MKG800, MIKX, China). Specifically, divergent primers annealing at the distal ends of circRNA were utilized to quantify circRNA abundance. Details of the primers can be found in **Supplementary Table 12**. Amplification was carried out using the StepOnePlus Real-Time PCR System (CFX, Bio-rad, USA), and Ct thresholds were determined by the software.

### Zebrafish gene editing based on CRISPR/Cas9 system

The knockout targets of zebrafish *sws1* and *sws2* genes were designed on the website of The University of California Santa Cruz (UCSC) Genome Browser (https://genome.ucsc.edu/). Synthetic PCR system of sgRNA: 2×TaqPCR Master Mix 10ul, target 1.5 μl, Oligo2 1.5 μl, ddH2O 7 μl. PCR procedure: 98 [for 2 min; 50 [for 10 min; 72 [for 10 min; Store at 4 [. The HiScribe T7 High Yield RNA Synthesis Kit (E2050, NEB, USA) was transcribed in vitro, and the gRNA was purified using the RNA Clean & Concentrator Kit (R1070, Zymo Research, USA). The synthesis experiment of Cas9 mRNA was purified by T3 plasmid pT3TS-nCas9n,

*Xba*[(R1045S, NEB) water bath for 2 h at 37 [, and PCR Purification Kit was used. The kit mMESSAGE mMACHINE™ T3 (AM1348, Thermo Fisher, USA) was used for *in vitro* transcription. Cas9 mRNA was obtained by purifying the transcripts with the RNA Clean & Concentrator Kit. The final concentrations of sgRNA and Cas9 mRNA were adjusted by injection system to 30 ng/μl and 50 ng/μl, and the mixed sgRNA and Cas9 mRNA were injected into the 1-cell stage embryos of wild type AB strain zebrafish by microinjection. DNA from zebrafish embryos 48 h after injection was extracted by NaOH alkaline cleavage method and amplified by PCR. PCR system: 2×Taq PCR Master Mix 10μL, DNA 3 μl, F+R 1 μl, ddH2O 6 μl. PCR procedure: 95 [5 min, 95 [30 sec, 60 [30 sec, 72 [30 sec, 35 cycles; 72 [for 10 min; Store at 4 [. The knock-out efficiency of PCR products was measured by Sanger sequencing.

The microinjected zebrafish embryos were cultured to 3-month-old adult fish, and F0 generation heterozygous mutants were screened by Sanger sequencing. Heterozygote and wild-type zebrafish were used to obtain F1 zebrafish, homozygous mutants were screened by Sanger sequencing and sent to Shanghai Sangong Biological Company for sequencing^72^.

### Behavioral testing

The behavioral test used to evaluate sws1 and sws2 knockout in zebrafish was the Y-maze experiment (20cm*50cm arm length), with the three arms of the Y-maze set as the starting arm, the light arm, and the dark arm, respectively, with equal amounts of feed in the light arm and the dark arm. 30 wild-type zebrafish and 30 mutant zebrafish were tested successively. Starting from the starting arm, each fish was tested for 2 minutes, and the final swimming direction was recorded. Each fish was tested three times.

### Vector construction and transfection

To construct opsin overexpression plasmid, we cloned human *sws*, zebrafish *sws1*, and *sws2* cDNAs and cloned them into pmcherry-N1 Vector using the Homologous Recombinant Kit (C113, Vazyme) (Supplementary Fig.7A). Amplification primers are shown in **Supplementary Table 12**. Transfection was performed using Lipo8000 (C0533FT, Beyotime, USA) according to the manufacturer’s instructions. After transfection, fluorescence expression was observed by fluorescence microscopy (IX53, Olympus, Japan), and untransfected HEK293 cells were used as controls.

### SA-**β**-Gal staining

SA-β-Gal staining was conducted utilizing the SA-β-Gal Staining Kit (G1073, Servicebio, China) following the manufacturer’s instructions. Synovial cells were fixed with senescent cell staining fixative for 15 min, washed 3 times and then incubated in SA-β-Gal staining solution at 37 [for 24 h. Images were captured under an inverted fluorescence microscope. SA-β-Gal positive synovial cells were randomly divided into three regions.

### Statistics

All the data from the study were expressed as Mean[±[SD. Analyses were performed using Prism 9.1.2 (GraphPad Software Inc.). The mean of the groups was compared using a Student’s t-test and ANOVA. Kaplan-Meier survival curves for mice and P-values were calculated using a log-rank test. P values of[<[0.05 indicate statistical significance.

### Data Availability Statement

The genome assembly, annotation files and all the sequencing data of the *Okamejei kenojei* have been deposited at China National GeneBank (CNGB) database^73^ with accession number CNP0003537, also at NCBI with BioProject number PRJNA990959, The *Okamejei kenojei* Genome Shotqun project has also been deposited at DDBJ/ENA/GenBank under the accession JAVHYY000000000.1. The GenBank assembly version described in this paper is GCA_035221665.1, as well as the genome assembly, annotation files and all the sequencing data of *Prionace glauca* at NCBI with BioProject number PRJNA1018302 and BioSample number SUB13848110. The *Prionace glauca* Genome Shotqun project has also been deposited at DDBJ/ENA/GenBank under the accession JBBMRE000000000. The GenBank assembly version described in this paper is GCA_037974335.1.

## Author Contributions

B.Z. and Y.L. initiated, managed, and drove genome sequencing project; B.B., B.Z., N.Z., Y.Z. and J.S. conceived the study; B.B, B.Z, and Y.L. designed the analysis; B.Z. and X.X. prepared the samples; M.L., L.J. and X.X performed the genome assembly and annotation; M.L. and L.J. performed the gene family and positive selection analysis; Y.F. and Y.L. performed the experiments and data analysis; B.B., B.Z., Y.L., L.J., Y.F, N.Z., M.L., X.X. and A.M. discussed the data. All authors contributed to data interpretation, B.B., B.Z., M.L., Y.F., Y.L. and N.Z. wrote and revised the paper with significant contributions from all other authors.

## Supplementary Material

Supplementary Material is available in the online version of the paper.

---

**Supplementary Fig.1:**
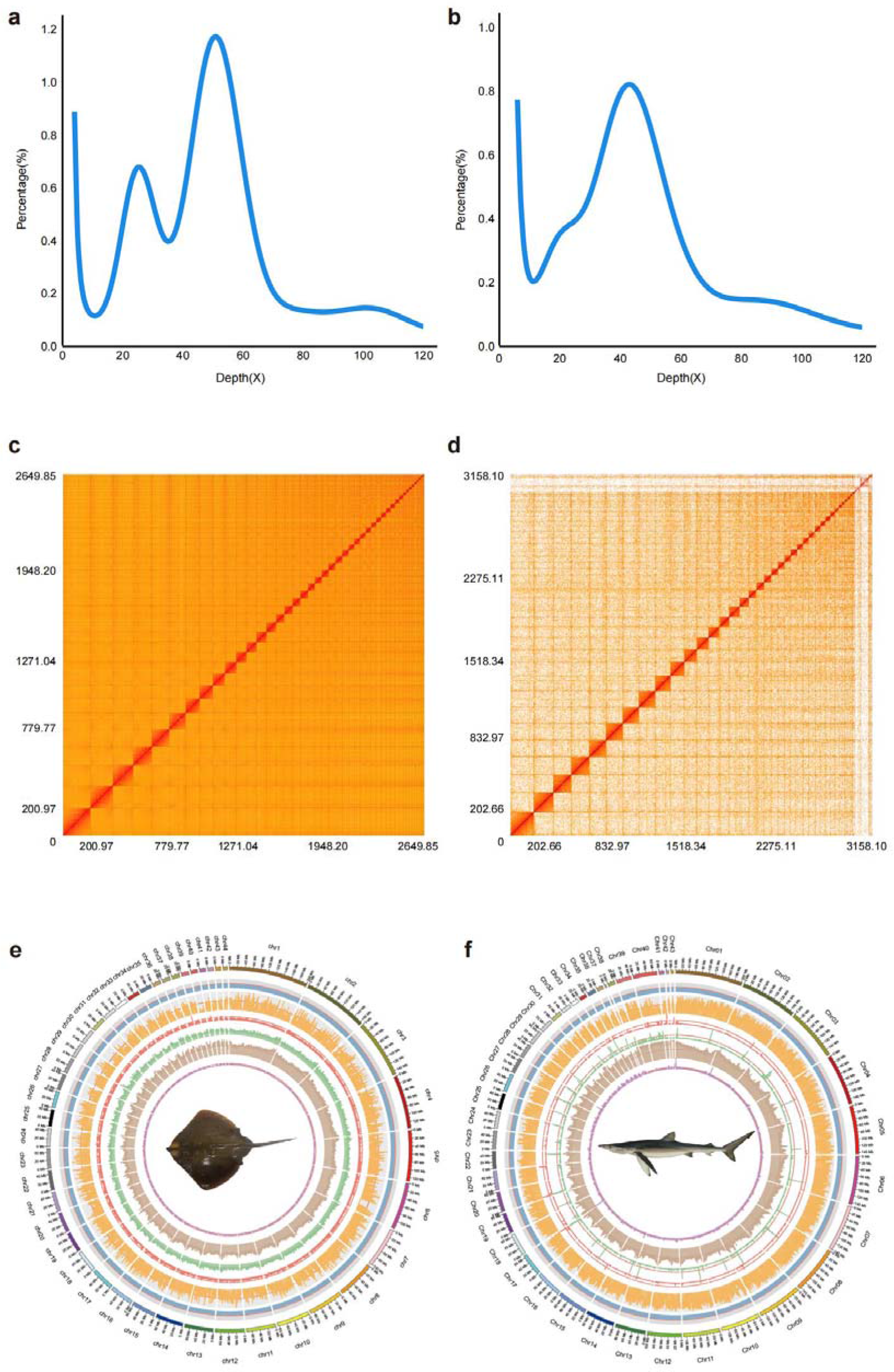
Characteristics of *O. kenojei* and *P. glauca* genomes. a,b K-mer frequency distribution of *O. kenojei* and *P. glauca* genomes, respectively (k=17). The estimated genome sizes of these two species are 2.79 and 3.25 Gb, respectively. c,d 44 and 43 chromosomes Hi-C contact maps of the *O. kenojei* and *P. glauca* genomes, respectively. e,f Circos plots of genome features of *O. kenojei* and *P. glauca*. From outer to inner circles: 1, represents chromosomes; 2, GC content; 3, gene density distribution; 4-7, distribution of DNA transposons, LTR, LINE and SINE. 2-7 are drawn with 500 Kb sliding windows.

**Supplementary Fig.2:**
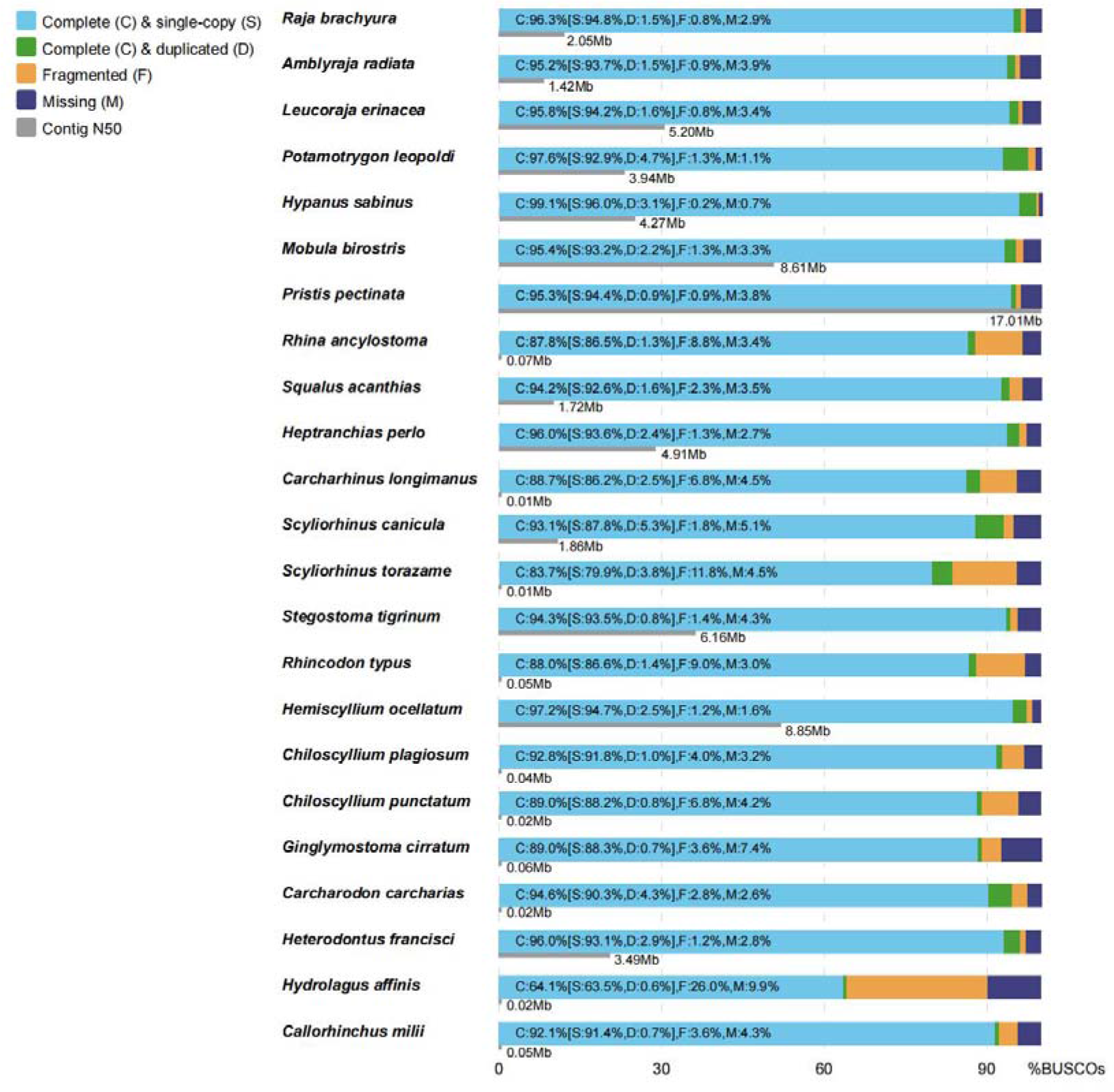
Comparison of BUSCOs and contig N50 of cartilaginous fish genomes. Comparison of BUSCOs (blue, green, orange and purple bars) and contig N50 (grey bar) of cartilaginous fish genomes.

**Supplementary Fig. 3.**
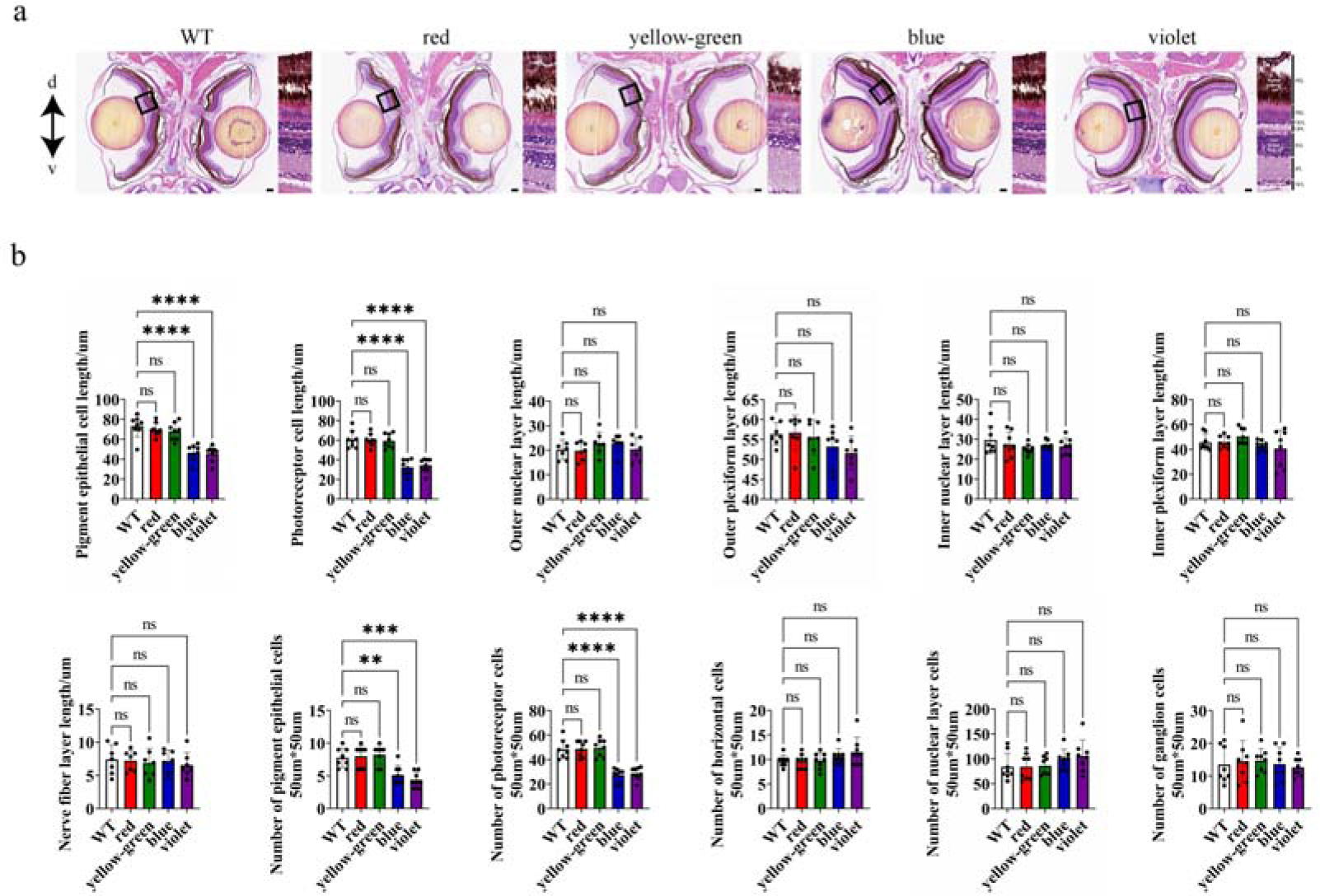
Shortwave light induced PEL and PRL damage indicated by histology presentation and statistics a. The picture shows the largest cross-section of the head of zebrafish after red, yellow-green, blue and violet light irradition. The zoomed-in view of the retina marked by black pane is displayed beside each panel. d, dorsal; v, ventral. Scale bar: 50um.b. The statistical results of pigment epithelial cells, photoreceptor cells, outer nuclear layer cells, horizontal cells, ganglion cells, the length of the pigment epithelial layer (PEL), photoreceptor cell layer(PRL), outer nuclear layer (ONL), outer plexus layer(OPL), inner nuclear layer(INL), inner plexus layer(IPL), and nerve fiber layer (NFL).Each bar in the histogram represents the average value, and the points represent the specific numerical distribution for each sample. n=7.**P<0.01,***P<0.001,****P<0.0001.

**Supplemental Fig. 4.**
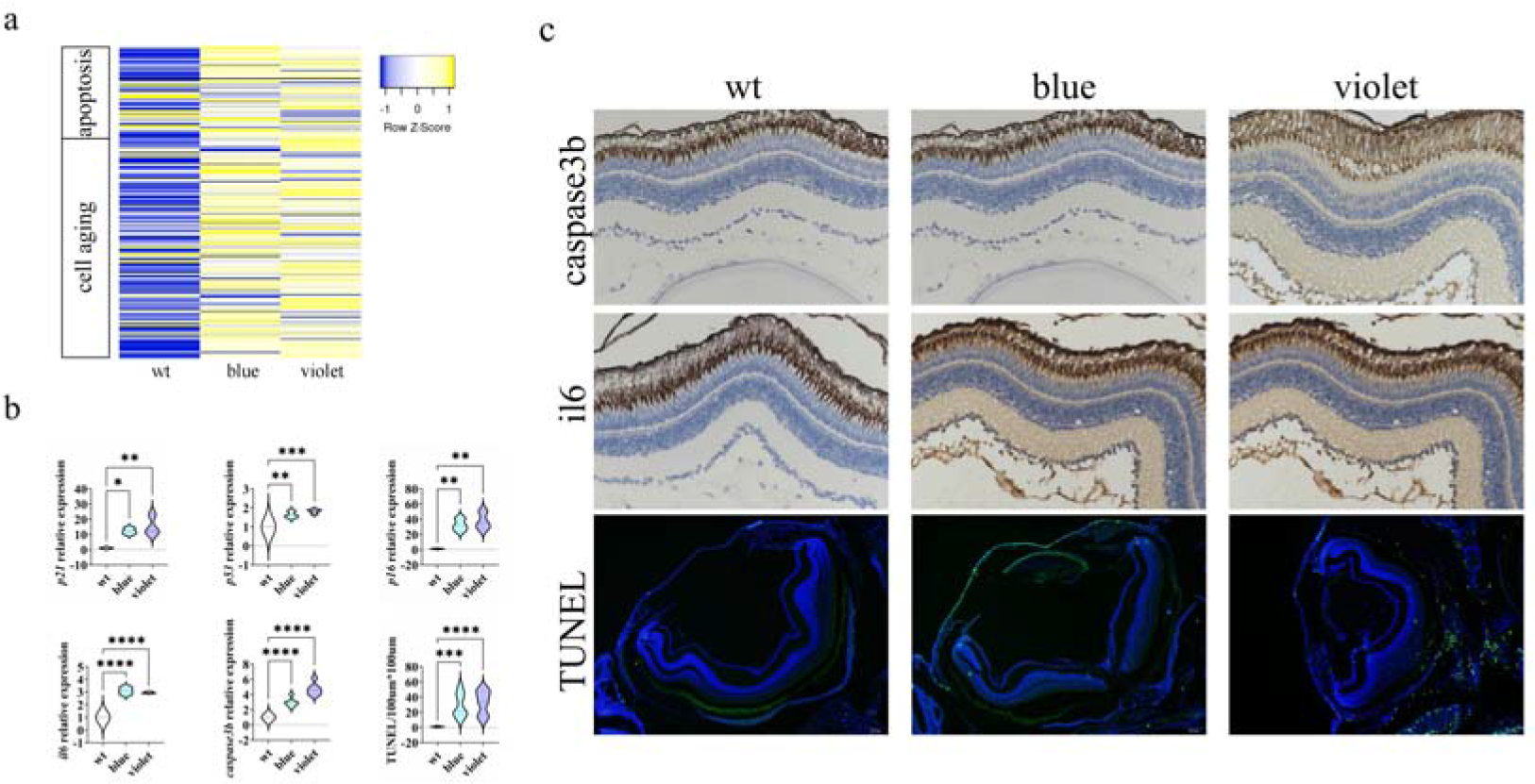
transcriptomic and immunostaining analysis demonstrated that Short-wave light caused both retinal apoptosis and cell aging. a. Comparison of Heat map from transcriptomic apoptosis-related genes and age-related genes in wild type of zebrafish between the white, blue and violet light irradiation. A total of 83 apoptosis-related genes were enriched, of which 54 genes were upregulated after shortwave light irradiation, and 198 cell-aging-related genes were upregulated after shortwave light irradiation. b. RT-qPCR of marker genes of the cell apoptosis (*p53*,*p21*,*caspase3b*) and cell aging (*p16*,*il6*), the internal reference gene is *b-actin*. In addition, the statistics on the number of DNA-breaking cells with TUNEL. n=3. *P<0.05, **P<0.01, ****P<0.0001. c. immunostaining of the apoptosis marker caspase3b and cell aging marker gene *il6*, and TUNEL fluorescence staining in zebrafish retina after light irradiation.

**Supplemental Fig. 5.**
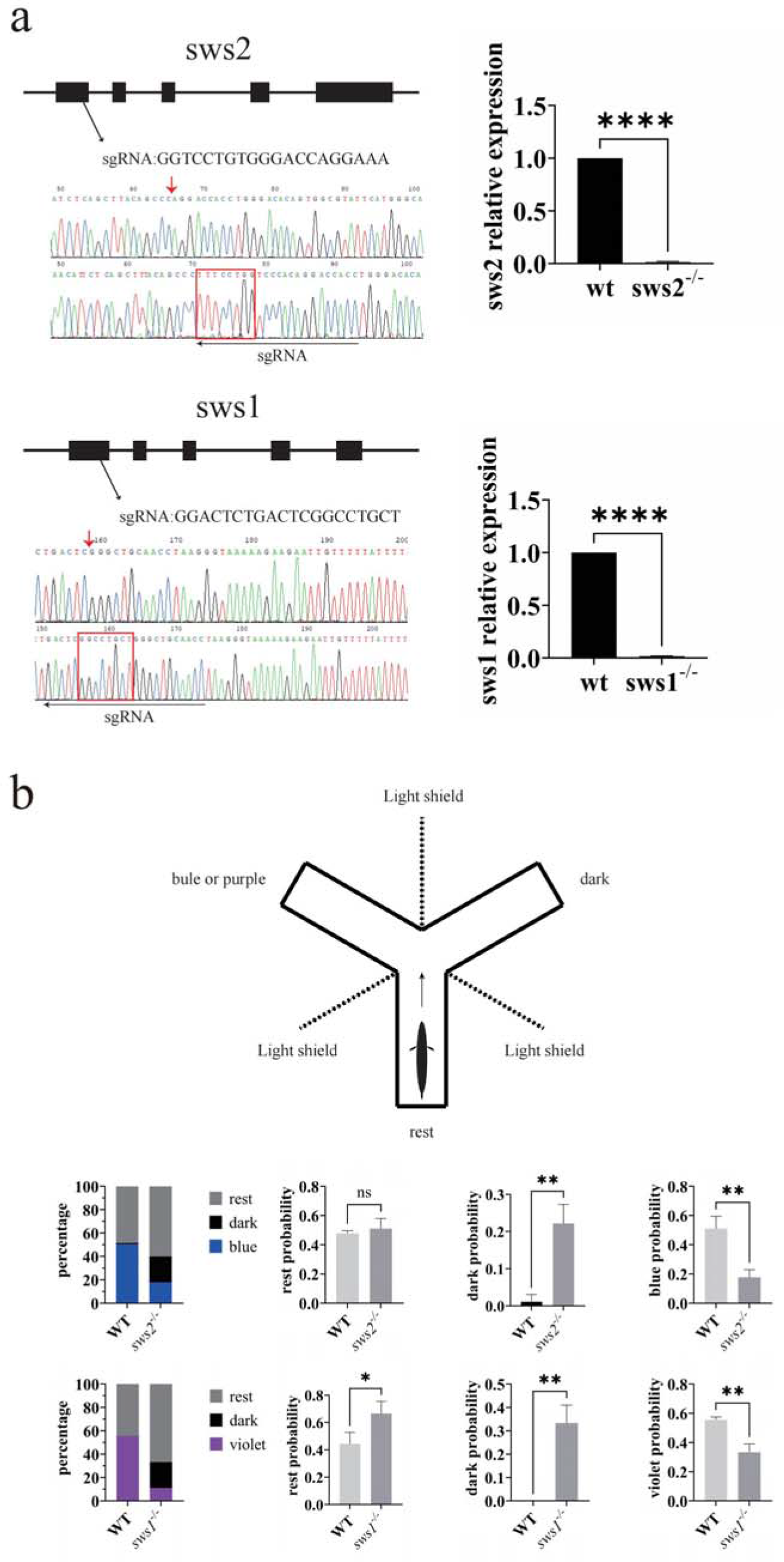
*sws2* and *sws1* knockouted respectively in zebrafish by CRISPR/cas9 technology. a. The black line segment indicates the sgRNA binding site and sequence, and the red arrow marks the origin site where the fragment is deleted. The red box highlights specific sequences that 8bp are deleted in sws2-/- and 14bp are deleted in sws1-/- . Primers were designed at the deletion site and the relative expression of sws was detected by qpcr. n=3.****P<0.0001. b. Schematic diagram of Y-maze experiment design and statistical results. The percentage of zebrafish swimming direction in each group was calculated, and the difference of three final swimming direction probabilities between wild type and mutant zebrafish was analyzed statistically.n=30.*P<0.05.**P<0.01.

**Supplemental Fig. 6:**
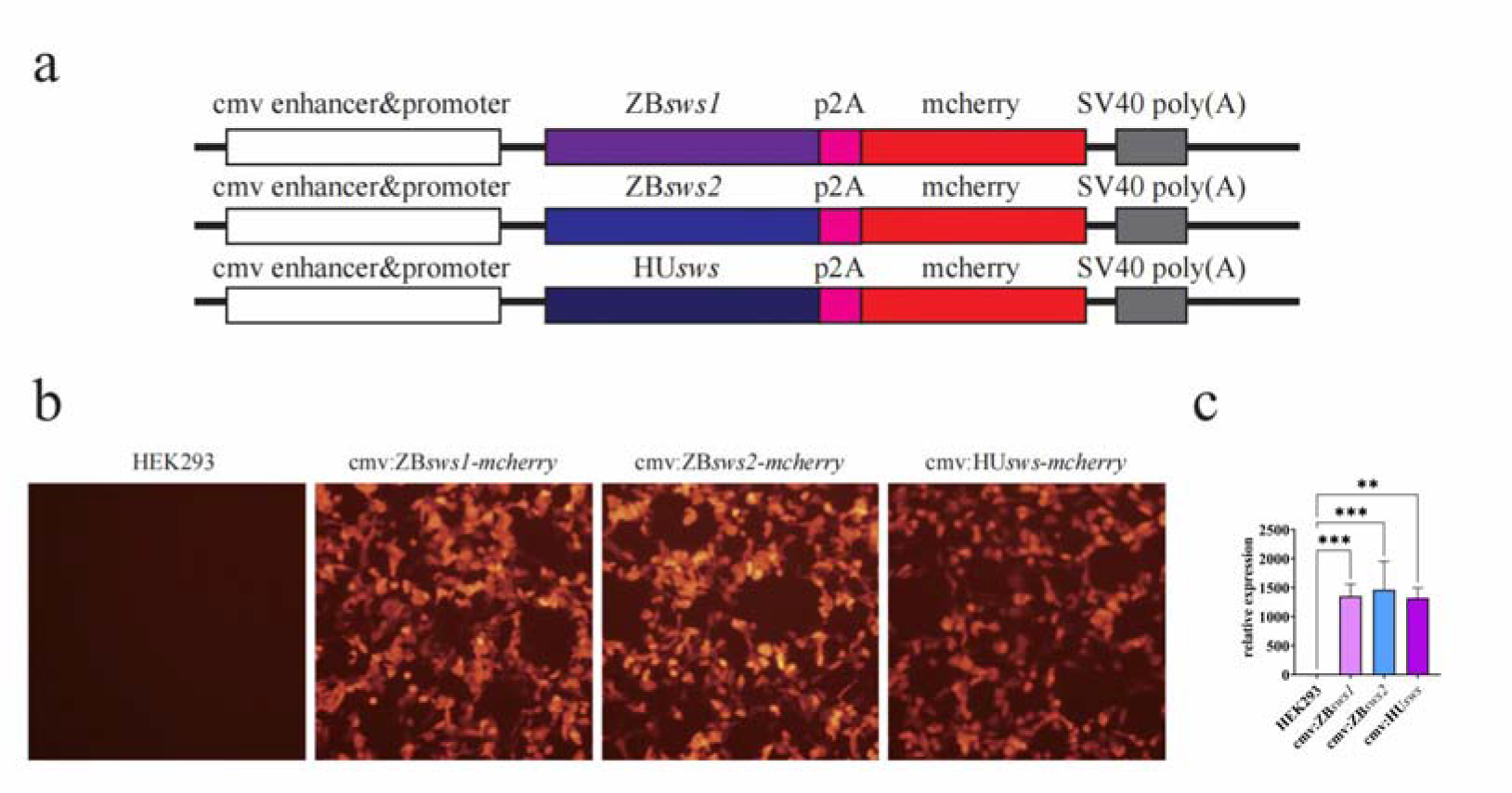
The construction of *sws*-overexpression cell vector a. Principle diagram of construction of *sws-*overexpression cell vector. ZB: zebrafish, HU: human, p2A: tupturing peptide, mcherry: red fluorescent protein, SV40 poly(A): transcription termination signal. b. The fluorescence results 48h after transfection of three plasmids into HEK293 cells. c. the relative expression of *sws* in cells. n=3. **P<0.01,***P<0.001.

