## Supplementary Table for "Adaptive loss of shortwave cone opsins during genomic evolution in cartilaginous fish"

Supplementary Tables

**Supplementary Table 1. Summary of the sequencing data.**

| ***O. kenojei*** | **Library** | | **Insert size (bp)** | | **Read length (bp)** | | **Bases (Gb)** | |
| --- | --- | --- | --- | --- | --- | --- | --- | --- |
|  | WGS  Hi-C | | 350 | | 100 | | 173.28 | |
|  |  |  | - | | 100 | | 473.83 | |
|  | **Library** | | **Reads number** | | **Length (Gb)** | | **Average length (bp)** | |
|  | Pacbio | | 15,876,797 | | 232.59 | | 14,649.63 | |
| ***P.glauca*** | | **Library** | | **Insert size (bp)** | | **Read length (bp)** | | **Bases (Gb)** |
|  | | WGS  Hi-C | | 350 | | 150 | | 156.98 |
|  | |  |  | - | | 150 | | 323.41 |
|  | | **Library** | | **Reads number** | | **Length (Gb)** | | **Reads LenN50 (bp)** |
|  | | CCS | | 5,057,066 | | 91.22 | | 18,666 |

**Supplementary Table 2. Statistics of the assembly genome information.**

|  | ***O. kenojei*** | ***P.glauca*** |
| --- | --- | --- |
| **Statistical level** | **Contig** | **Contig** |
| Total number | 5,781 | 1,904 |
| Total length (bp) | 2,754,429,125 | 3,227,307,048 |
| Average length (bp) | 476,462 | 1,695,014 |
| N50 Length (bp) | 1,503,252 | 5,319,141 |
| N90 Length (bp) | 204,937 | 657,817 |
| GC content (%) | 43.90 | 44.23 |

**Supplementary Table 3. BUSCO validation of the genome and geneset.**

|  | **BUSCO** | **Total** | **Complete** | **Complete and single-copy** | **Complete and duplicated** | **Fragmented** | **Missing** |
| --- | --- | --- | --- | --- | --- | --- | --- |
| *O. kenojei* | Genome | 954 | 895 (93.9%) | 884  (92.7%) | 11  (1.2%) | 16  (1.7%) | 43  (4.4%) |
|  | Gene | 954 | 853 (89.4%) | 837  (87.7%) | 16  (1.7%) | 41  (4.3%) | 60  (6.3%) |
| *P.glauca* | Genome | 954 | 877 (91.93%) | 868 (90.99%) | 9  (0.94%) | 22  (2.31%) | 55  (5.77%) |
|  | Gene | 954 | 873 (91.5%) | 855  (89.6%) | 18  (1.9%) | 8  (0.8%) | 73  (7.7%) |

**Supplementary Table 4. Function annotation of the genes in *O.kenojei* and *P.glauca* genome.**

|  | **Database** | **Number** | **Percent (%)** |
| --- | --- | --- | --- |
| *O. kenojei* | Total | 22,965 | 100% |
|  | Swiss-Prot | 19,734 | 85.93% |
|  | KEGG | 22,227 | 96.79% |
|  | TrEMBL | 22,047 | 96.00% |
|  | Interpro | 20,543 | 89.45% |
|  | Overall | 22,279 | 97.01% |
|  | **Database** | **Number** | **Percent (%)** |
| *P.glauca* | Total | 21,229 | 100% |
|  | Swiss-Prot | 18,550 | 87.38% |
|  | KEGG | 20,581 | 96.95% |
|  | TrEMBL | 20,981 | 98.83% |
|  | Interpro | 19,781 | 93.18% |
|  | Overall | 20,989 | 98.87% |

**Supplementary Table 5. Statistics of transposable elements of the *O.kenojei* genome.**

| **Type** | **RepBase TEs** | | **TE Proteins** | | ***De novo*** | | **Combined TEs** | |
| --- | --- | --- | --- | --- | --- | --- | --- | --- |
|  | **Length (bp)** | **% in genome** | **Length (bp)** | **% in genome** | **Length (bp)** | **% in genome** | **Length (bp)** | **% in genome** |
| DNA | 78,464,116 | 2.85 | 18,727,923 | 0.68 | 173,736,886 | 6.31 | 230,347,777 | 8.36 |
| LINE | 573,956,132 | 20.84 | 585,963,394 | 21.27 | 1,344,029,244 | 48.80 | 1,517,115,565 | 55.08 |
| SINE | 111,952,940 | 4.06 | 0 | 0.00 | 3,189,261 | 0.12 | 114,816,091 | 4.17 |
| LTR | 103,428,943 | 3.76 | 109,520,713 | 3.98 | 581,162,426 | 21.10 | 661,425,266 | 24.01 |
| Other | 9,421 | 0.00 | 0 | 0.00 | 0 | 0.00 | 9,421 | 0.00 |
| Unknown | 0 | 0.00 | 0 | 0.00 | 366,054 | 0.01 | 366,054 | 0.01 |
| Total | 837,145,201 | 30.39 | 713,843,693 | 25.92 | 1,856,577,927 | 67.40 | 1,908,097,922 | 69.27 |

**Supplementary Table 6. Statistics of transposable elements of the *P.glauca* genome.**

| **Type** | **RepBase TEs** | | **TE Proteins** | | ***De novo*** | | **Combined TEs** | |
| --- | --- | --- | --- | --- | --- | --- | --- | --- |
|  | **Length (bp)** | **% in genome** | **Length (bp)** | **% in genome** | **Length (bp)** | **% in genome** | **Length (bp)** | **% in genome** |
| DNA | 64,540,161 | 2.00 | 119,059 | 0.00 | 6,796,243 | 0.21 | 70,701,830 | 2.19 |
| LINE | 803,476,156 | 24.90 | 810,490,848 | 25.11 | 2,022,660,369 | 62.67 | 2,064,840,715 | 63.98 |
| SINE | 223,363,271 | 6.92 | 0 | 0.00 | 7,322,807 | 0.23 | 227,879,434 | 7.06 |
| LTR | 62,679,115 | 1.94 | 8,666,941 | 0.27 | 85,662,804 | 2.65 | 142,002,652 | 4.40 |
| Other | 3,891 | 0.00 | 0 | 0.00 | 1,942 | 0.00 | 4,372 | 0.00 |
| Unknown | 0 | 0.00 | 0 | 0.00 | 2,337,375 | 0.07 | 2,337,375 | 0.07 |
| Total | 1,113,942,257 | 34.51 | 819,202,795 | 25.38 | 2,092,898,793 | 64.85 | 2,144,308,681 | 66.44 |

**Supplementary Table 7. Summary of the RNA sequencing data of *O. kenojei*.**

| **Sample** | **Tissue** | **Total Bases (Gb)** |
| --- | --- | --- |
| TQD200704374-1-C-Ey-1 | Eye | 13.99 |
| TQD200704375-1-C-Ou-1 | Oarium | 15.43 |
| TQD200704377-1-C-SKB-1 | Dorsal skin | 11.41 |
| TQD200704380-1-C-Li-1 | Liver | 12.88 |
| TQD200704381-1-C-SKF-1 | Abdomen skin | 13.52 |
| TQD200704383-1-C-Pt-1 | Spermary | 19.05 |
| TQD200704386-2-X-Pt-1 | Spermary | 16.24 |
| TQD200704396-2-X-Ou-1 | Oarium | 12.91 |
| Total | - | 115.44 |

**Supplementary Table 8. Published genomes used in repeat analysis.**

| **Species** | **Accession** |
| --- | --- |
| *Raja brachyura* | GCA_963514005.1 |
| *Amblyraja radiata* | GCF_010909765.2 |
| *Leucoraja erinacea* | GCF_028641065.1 |
| *Potamotrygon leopoldi* | Article^1^ |
| *Pristis pectinata* | GCA_009764475.2 |
| *Hypanus sabinus* | GCF_030144855.1 |
| *Mobula birostris* | GCA_030035685.1 |
| *Squalus acanthias* | GCA_030390025.1 |
| *Stegostoma tigrinum* | GCF_030684315.1 |
| *Rhincodon typus* | GCF_021869965.1 |
| *Chiloscyllium plagiosum* | GCF_004010195.1 |
| *Hemiscyllium ocellatum* | GCF_020745735.1 |
| *Scyliorhinus canicula* | GCF_902713615.1 |
| *Carcharodon carcharias* | GCA_003604245.1 |
| *Scyliorhinus torazame* | GCA_003427355.1 |
| *Chiloscyllium punctatum* | GCA_003427335.1 |
| *Callorhinchus milii* | GCF_000165045.1 |

**Supplementary Table 9. Published protein sequences used in gene annotation.**

| **Species** | **Accession** |
| --- | --- |
| *Chiloscyllium punctatum* | GCA_003427335.1 |
| *Scyliorhinus torazame* | GCA_003427355.1 |
| *Callorhinchus milii* | GCF_000165045.1 |
| *Rhincodon typus* | GCF_001642345.1 |
| *Carcharodon carcharias* | Article^2^ |

**Supplementary Table 10. Published protein sequences used in gene family analysis.**

| **Species** | **Accession** |
| --- | --- |
| *Amblyraja radiata* | GCF_010909765.2 |
| *Chiloscyllium plagiosum* | GCF_004010195.1 |
| *Callorhinchus milii* | GCF_000165045.1 |
| *Carcharodon carcharias* | Article^2^ |
| *Chiloscyllium punctatum* | GCA_003427335.1 |
| *Rhincodon typus* | GCF_001642345.1 |
| *Scyliorhinus torazame* | GCA_003427355.1 |
| *Latimeria chalumnae* | GCF_000225785.1 |
| *Lepisosteus oculatus* | GCF_000242695.1 |
| *Danio rerio* | GCF_000002035.6 |
| *Acipenser ruthenus* | GCF_010645085.1 |
| *Coilia nasus* | Article^3^ |
| *Gasterosteus aculeatus* | GCF_016920845.1 |
| *Takifugu rubripes* | GCF_901000725.2 |
| *Scleropages formosus* | GCF_900964775.1 |

**Supplementary Table 11. Published protein sequences to identify opsin genes.**

| **Species** | **Accession** |
| --- | --- |
| *Amblyraja radiata* | GCF_010909765.2 |
| *Leucoraja erinacea* | GCF_028641065.1 |
| *Potamotrygon leopoldi* | Article^1^ |
| *Hypanus sabinus* | GCF_030144855.1 |
| *Pristis pectinata* | GCF_009764475.1 |
| *Stegostoma tigrinum* | GCF_030684315.1 |
| *Rhincodon typus* | Article^4^ |
| *Chiloscyllium plagiosum* | GCF_004010195.1 |
| *Chiloscyllium punctatum* | Article^4^ |
| *Hemiscyllium ocellatum* | GCF_020745735.1 |
| *Scyliorhinus canicula* | GCF_902713615.1 |
| *Scyliorhinus torazame* | Article^4^ |
| *Carcharodon carcharias* | GCF_017639515.1 |
| *Callorhinchus milii* | GCF_000165045.1 |
| *Lepisosteus_oculatus* | GCF_000242695.1 |
| *Takifugu rubripes* | GCF_901000725.2 |
| *Gasterosteus aculeatus* | GCF_016920845.1 |
| *Danio rerio* | GCF_000002035.6 |
| *Clupea harengus* | GCF_900700415.2 |
| *Sardina pilchardus* | Article^5^ |
| *Coilia nasus* | Article^3^ |
| *Scleropages formosus* | GCF_900964775.1 |
| *Acipenser ruthenus* | GCF_010645085.1 |
| *Latimeria chalumnae* | GCF_000225785.1 |

**Supplementary Table 12. Primers used in the qRT-PCR of the study**

| gene | primers |
| --- | --- |
| β-actin-F | GGAAATCGTGCGTGACATTAAG |
| β-actin-R | CCTCTGGACAACGGAACCTCT |
| caspase3b-F | TCTTCGAGTTTGGTGGGACCATGT |
| caspase3b-R | TACATCTCCACGAAGGCATCCCAA |
| bcl2-F | TGAGGCTCTACCGGGTGTTA |
| bcl2-R | ACATGGTCCCACCAAACTCG |
| il6-F | ATGTCTAACGCGAATCTACAGC |
| il6-R | GTCTGATCCATCTCTCCGTCT |
| MMP65-F | TTTCTACACACCCACTGGCAAC |
| MMP65-R | ACTCCTTGCCATACACCGAT |
| p21-F | CGGCTTTCTGAAGTTTAGCAT |
| p21-R | CTGTGCAAGCTACACTACTCC |
| P53-F | TGCTACTAAACTACATGTGCAA |
| P53-R | GCACCACATCACTTAACTCC |
| sws1-target | ggactctgactcggcctgct |
| Sws2-target | ggtcctgtgggaccaggaaa |
| sws1-F | cacgcaggagctctttgaga |
| sws1-R | cgttttagatctgtgcgtcca |
| Sws2-F | aacagcaaacgccagaac |
| Sws2-R | ctaccgaggaaccgaaaa |
| Oligo2 | AAAAGCACCGACTCGGTGCCACTTTTTCAAGTTGATAACGGACTAGCCTTATTTTAACTTGCTATTTCTAGCTCTAAAAC |

1	Zhou, J. *et al.* Draft Genome of White-blotched River Stingray Provides Novel Clues for Niche Adaptation and Skeleton Formation. *Genomics, proteomics & bioinformatics* **21**, 501-514, doi:10.1016/j.gpb.2022.11.005 (2023).

2	Marra, N. J. *et al.* White shark genome reveals ancient elasmobranch adaptations associated with wound healing and the maintenance of genome stability. *Proceedings of the National Academy of Sciences of the United States of America* **116**, 4446-4455, doi:10.1073/pnas.1819778116 (2019).

3	Xu, G. *et al.* Genome and population sequencing of a chromosome-level genome assembly of the Chinese tapertail anchovy (Coilia nasus) provides novel insights into migratory adaptation. *GigaScience* **9**, doi:10.1093/gigascience/giz157 (2020).

4	Hara, Y., Yamaguchi, K. & Onimaru, K. Shark genomes provide insights into elasmobranch evolution and the origin of vertebrates. **2**, 1761-1771, doi:10.1038/s41559-018-0673-5 (2018).

5	Louro, B. *et al.* A haplotype-resolved draft genome of the European sardine (Sardina pilchardus). *GigaScience* **8**, doi:10.1093/gigascience/giz059 (2019).
